# A multiscale model of striatum microcircuit dynamics

**DOI:** 10.1101/2023.12.28.573546

**Authors:** Federico Tesler, Alexander Kozlov, Sten Grillner, Alain Destexhe

**Affiliations:** Paris-Saclay University, CNRS, Paris-Saclay Institute of Neuroscience (NeuroPSI), 91198 Gif-sur-Yvette, France; Science for Life Laboratory, School of Electrical Engeneering and Computer Science, Royal Institute of Technology, SE-10044 Stockholm, Sweden; Department of Neuroscience, Karolinska Institutet, SE-17165 Stockholm

## Abstract

The striatum is the largest structure in the basal ganglia, and is known for its key role in functions such as learning and motor control. Studying these aspects requires investigating cellular/microcircuits mechanisms, in particular related to learning, and how these small-scale mechanisms affect large-scale behavior, and its interactions with other structures, such as the cerebral cortex. In this paper, we provide a multiscale approach to investigate these aspects. We first investigate striatum dynamics using spiking networks, and derive a mean-field model that captures these dynamics. We start with a brief introduction to the microcircuit of the striatum and we describe, step by step, the construction of a spiking network model, and its mean-field, for this area. The models include explicitly the different cell types and their intrinsic electrophysiological properties, and the synaptic receptors implicated in their recurrent interactions. Then we test the mean-field model by analyzing the response of the striatum network to the main brain rhythms observed experimentally, and compare this response to that predicted by the mean-field. We next study the effects of dopamine, a key neuromodulator in the basal ganglia, on striatal neurons. Integrating dopamine receptors in the spiking network model leads to emerging dynamics, which are also seen in the mean-field model. Finally, we introduce a basic implementation of reinforcement learning (one of the main known functions of the basal-ganglia) using the mean-field model of the striatum microcircuit. In conclusion, we provide a multiscale study of the striatum microcircuits and mean-field, that capture its response to periodic inputs, the effect of dopamine and can be used in reinforcement learning paradigms. Given that several mean-field models have been previously proposed for the cerebral cortex, the mean-field model presented here should be a key tool to investigate large-scale interactions between basal ganglia and cerebral cortex, for example in motor learning paradigms, and to integrate it in large scale and whole-brain simulations.

## 1 Introduction

The striatum is the largest structure in the basal ganglia and it is known for its key role in multiple brain functions such as motor control, learning and decision making [1–3]. The important role played by the striatum emerges from the complex interplay between its cellular and structural composition and its interaction with other brain areas. Hence, understanding and modelling the complex dynamics and functions of the striatum constitutes a very relevant and challenging task. In terms of cellular composition, the majority of neurons in the stratium correspond to striatal projection neurons (SPN) which represent around 95% of the cells in rodents and 85% in primates, forming a dense network of lateral and recurrent synaptic connections [4, 5]. Regarding the interaction with other regions, the striatum receives major glutamatergic input from the cortex, in particular from the sensorimotor cortex and associative regions of cortex. In addition, it receives input from the thalamus and other basal-ganglia nuclei, being particularly relevant the dopaminergic projections from substantia nigra pars compacta (SNc) [5, 6]. In turn the striatum projects back to these different regions and nuclei via two main pathways, known normally as the *direct* and *indirect* pathways: nearly half of the SPN cells project directly to the output nuclei of the basal ganglia (internal globus palidus, GPi, and substantia nigra pars reticulata, SNr) which project themselves to the thalamus and downstream motor centers. This type of SPN cells are referred to as *direct pathway projection neurons* (dSPN). The other half of SPN cells projects to the external globus pallidus (GPe), which in turn projects to the subthalamic nucleus and output nuclei, and they are referred to as *indirect pathway striatal projection neurons* (iSPN) [4–6] (see Fig. 1.a for a diagram of mouse basal ganglia). The dSPN are known to initiate movements through disinhibition of brainstem motor centers, while the net effect of iSPN cells is to further enhance the inhibitory activity of the basal ganglia output nuclei (inhibiting motion) [4].

**Fig. 1.**
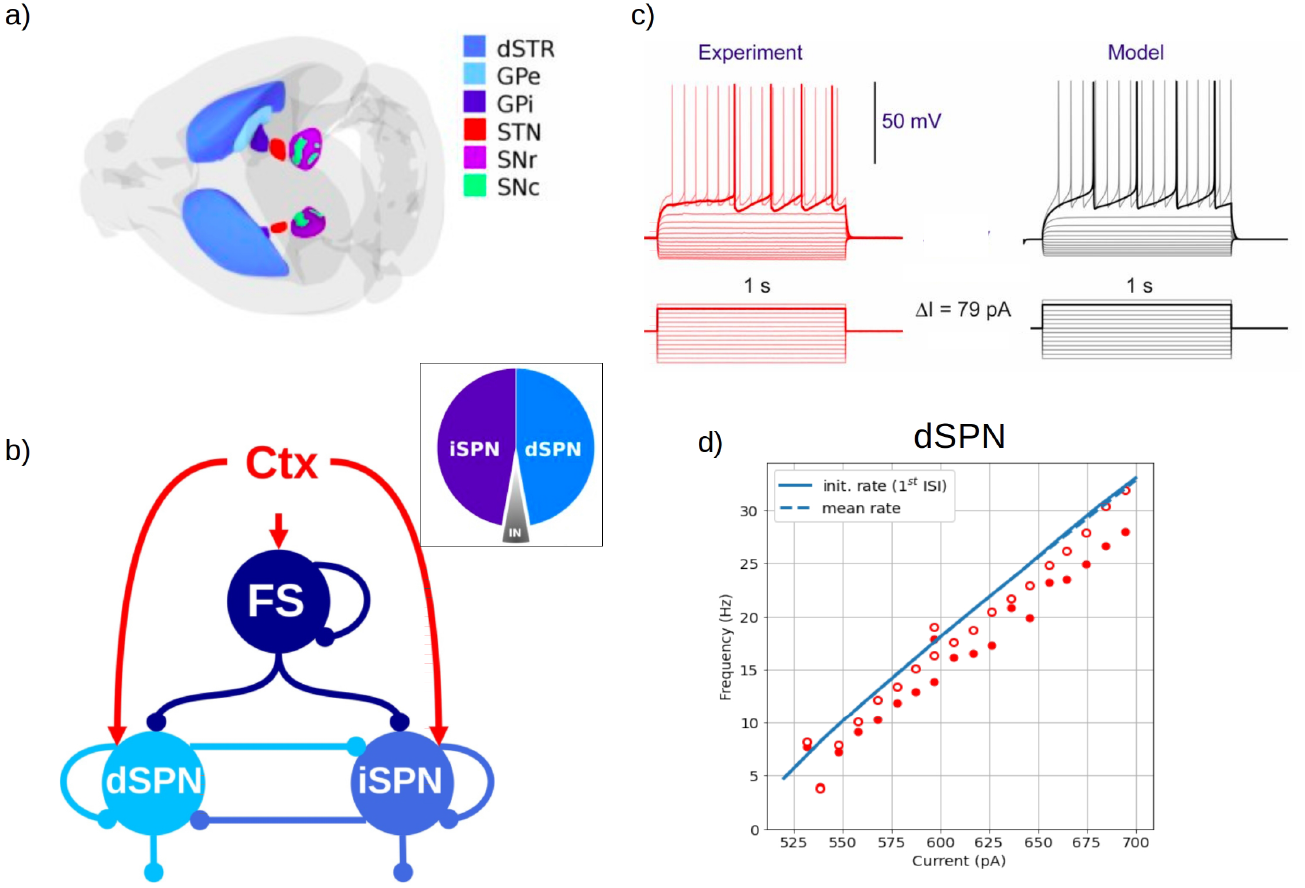
a) Dorsal view of the mouse brain showing the basal ganglia subnuclei. The dorsal striatum (dSTR), globus pallidus external and internal segment (GPe and GPi, respectively), subthalamic nucleus (STN), substantia nigra pars reticulata and pars compacta (SNr and SNc, respectively) are shown in relative sizes. b) Diagram of the connectivity within the striatum between direct and indirect SPN cells, FS cells and cortex (Ctx) projections to the striatum. Inset: relative distribution of each cell-type within the striatum. SPN cells correspond to about 95% of the total, while the rest corresponds to striatal interneurons. c) Example of the response of an dSPN cell to the injection of current of different amplitude for experiments (left) and for the AdEx neuronal model (right). d) Example of a fit of the AdEx model on experimental data for dSPN cells. Panels (a) and (c) adapted from Ref.[4].

Thus, modelling and studying the functions of the striatum and its associated neuronal dynamics requires to investigate these cellular/microcircuits mechanisms, and how the small-scale mechanisms affect large-scale behavior. To build a model across these multiple scales represents a challenge, as it normally requires a compromise between the level of detail of the model and the need of computational resources. A useful approach to study large-scale neuronal dynamics consists in the use of mean-field models to analyze the activity of large neuronal populations or brain areas. Mean-field models use statistical techniques to estimate the activity of large populations (from hundreds to thousands of neurons), which allows to reduce high-dimensional systems into low dimensions based on the first few statistical moments. However, most of the existing mean-fields are based on generic models (sometimes inspired by cortical circuits) [7, 8], which do not consider the rich and specific cellular and synaptic variability observed along brain regions. In addition, most of the existing mean-field models cannot be directly linked to molecular and cellular parameters, for which an accurate transition between scales is usually not possible. Only recently the firsts detailed meanfield models of specific sub-cortical microcircuits have been proposed [9–11]. Thus, it is currently of great relevance to develop mean-field models that take into account the specificities of each region and that can incorporate biologically relevant parameters at the different scales.

In this context, we present in this paper a multiscale model of the striatum for which we introduce a novel mean-field model of the striatum microcircuit dynamics. To develop this mean-field model we will make use of a recently introduced formalism that follows a bottom-up approach (starting from the single-cell level), which allows to build a mean-field model that incorporates different cell types with specific intrinsic firing properties, and their specific synaptic interactions mediated by different receptor types [12–14]. This approach permits an efficient transition between scales and, furthermore, it allows to explore the effects of cellular parameters at the network level, as we will show for the case of dopaminergic effects in the stratium.

In the next sections we will first present a spiking network model of the striatum microstructure and the strategy to construct a striatal mean-field model. Then we will test and validate our approach by analyzing the response of the mean-field under the main oscillatory activity observed in the striatum and comparing its results to the spiking network model. To analyze a first potential use of the model, we show how the effect of the dopaminergic input can be incorporated in our model, which is of great relevance also for the study of neurological disorders such as Parkinson’s disease and schizophrenia. We compare the results obtained from the mean-field with the ones of the spiking-network, which further validates our model. In addition it illustrates how changes at the cellular level can lead to emerging effects at the network level, which can be captured by the mean-field model. Finally, we present a basic implementation of Reinforcement Learning with the mean-field model, which shows the capabilities of the model to reproduce specific brain functions and in particular its potential implementation for modelling behavioural experiments.

## 2 Methods

### 2.1 Single-cell model

To perform spiking-network simulations we will make use of the Adaptive Exponential Integrate-and-Fire neuronal model (AdEx). The equations for the AdEx model are given by [15]:

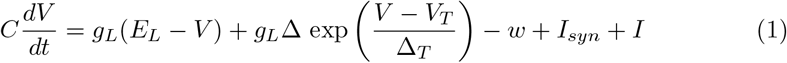

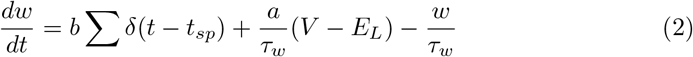

where *V* is the membrane potential, *w* is an adaptation current, *C* is the membrane capacity, *g*_*L*_ is the leakage conductance, *E*_*L*_ is the leakage reversal potential, *V*_*T*_ the threshold, Δ a slope factor, *I*_*syn*_ is the synaptic current, *I* represents any non-synaptic current acting on the cell, *a* is the sub-threshold adaptation constant and *b* is the spiking adaptation constant. When *V > V*_*T*_ at time *t* = *t*_*sp*_ a spike is generated, the membrane potential is reset to a voltage *V*_*res*_ and remains at this value for a refractory time *t*_*ref*_, and the adaptation variable is increased by an amount *b*. The synaptic current *I*_*syn*_ received by a neuron is given by the spiking activity of all presynaptic neurons connected to it. The total synaptic current can be written as a sum of the excitatory and inhibitory synaptic activity *I*_*syn*_ = Σ _*j*_ *G*_*j*_(*E*_*j*_ − *V*), where *j* correspond to each synaptic type, *E*_*j*_ is the synaptic reversal potential and *G*_*j*_ is the synaptic conductance. We model the synaptic conductances as decaying exponential functions that experience a quantal increase *Q*_*j*_ at each pre-synaptic spike: 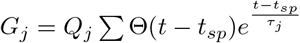, with *τ*_*j*_ the decay constant.

### 2.2 Striatum microcircuit

To study the neuronal dynamics of the striatal microcircuit to cortical input we will moodel a network made of direct striatal projection neurons (dSPN), indirect striatal projection neurons(iSPN) and Fast-Spiking interneurons (FS) (see diagrams in Fig.1.a,b, and Introduction) [4, 5]. Each cell type is modelled by the AdEx neuronal model described in the previous section. Single-cell parameters for each cell type were obtained by fitting on experimental data (Fig.1.c). Electrophysiological data correspond to mouae-striatum cells, presented in Hjorth et al, 2020, and available at https://kg.ebrains.eu/search/instances/Model/a6458de3-a176-4378-9b03-34a26f5da3bd. All cell types receive excitatory input from cortical projections. We consider an *Erdős–Rényi* network made of 1000 dSPN, 1000 iSPN and 100 FS cells. Based on previous work [4, 5], to preserve inhibitory synaptic input to individual cells, we set connection probability of FS cells to other FS cells *p*_*FS*−*FS*_ = 0.25 and to dSPN and iSPN cells *p*_*FS*−*dSPN*_ = 0.45 and *p*_*FS*−*iSPN*_ = 0.25, respectively. SPN cells project to other SPN cells in this setup with connectivity probability of *p*_*dSPN*−*dSPN*_ = 0.5, *p*_*dSPN*−*iSPN*_ = 0.1, *p*_*iSPN*−*iSPN*_ = 0.9 and *p*_*iSPN*−*dSPN*_ = 0.5, respectively.

### 2.3 Mean-Field formalism

We will adopt a recently developed Mean-Field formalism for AdEx neurons [13, 14]. The formalism is based on a bottom-up approach, starting at single-cell level, which allows the construction of mean-field models with cellular-type specificity. The meanfield equations for the AdEx network are given to a first-order by [14]:

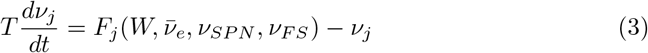

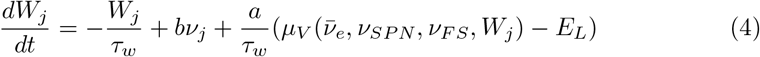

where *ν*_*j*_ is the mean neuronal firing rate of the population *j* (*j* =dSPN, iSPN, FS), *W*_*j*_ is the mean value of the adaptation variable for population *j, F* is the neuron transfer function (i.e. output firing rate of a neuron when receiving the corresponding excitatory and inhibitory inputs with mean rates 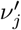 and with a level of adaptation W), *a* and *b* are the sub-threshold and spiking adaptation constants, *t*_*w*_ is the characteristic time of the adaptation variable and *T* is a characteristic time for neuronal response (we adopt *T* = 5 ms). The cortical input to the striatum is simulated as an (excitatory) external drive *ν*_*e*_.

Following Zerlaut et al (2018) [13] we write the transfer function for each neuronal type as:

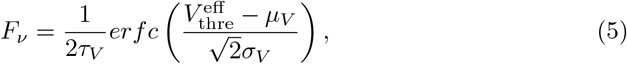

where *erfc* is the error function, 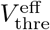 is an effective neuronal threshold, *µ*_*V*_, *σ*_*V*_ and *τ*_*V*_ are the mean, standard deviation and correlation decay time of the neuronal membrane potential. The effective threshold can be written as a second order polynomial expansion:

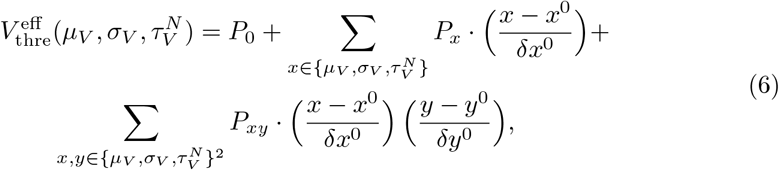

where *x*^0^, *y*^0^, *δx*^0^, *δy*^0^ are constants, and where the coefficients *P*_*xy*_ are to be determined by a fit over the numerical transfer function obtained from single-cell spiking simulations for each specific cell-type.

The mean membrane potential and standard deviation can be written as [13]:

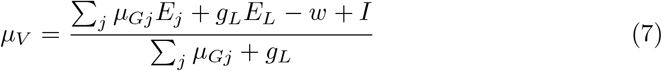

where 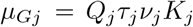 is the mean synaptic conductance and *K*_*j*_ is the mean synaptic convergence of type *j*.

Finally, we can write the standard deviation and correlation decay time of the neuronal membrane potential as [13]:

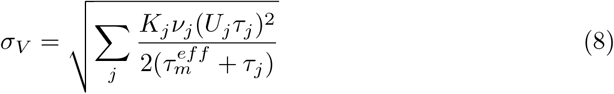

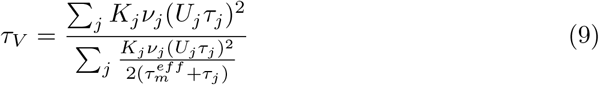

with 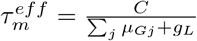 and 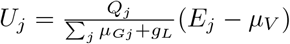

The detailed derivation of the mean field equations can be found in Refs. [13, 14].

## 3. Results

### 3.1 Mean-Field model of the Striatum Microcircuit

The first step in the semi-analytical approach to derive the mean-field model (see Methods) consists in the estimation of the transfer function for each cell-type (dSPN, iSPN, FS). This estimation is performed through the fit of the output-firing rate of each single-cell type with Eq.5, from where the parameters of the effective threshold *V*_*eff*_ are calculated (Eq.6). We show in Figure 2 the corresponding output firing rate and transfer function estimation for each cell-type. For illustration purposes in the figure we show the results of dSPN cells for *ν*_*FS*_ = 0, nevertheless the fit was performed for the full range in the three variables.

**Fig. 2.**
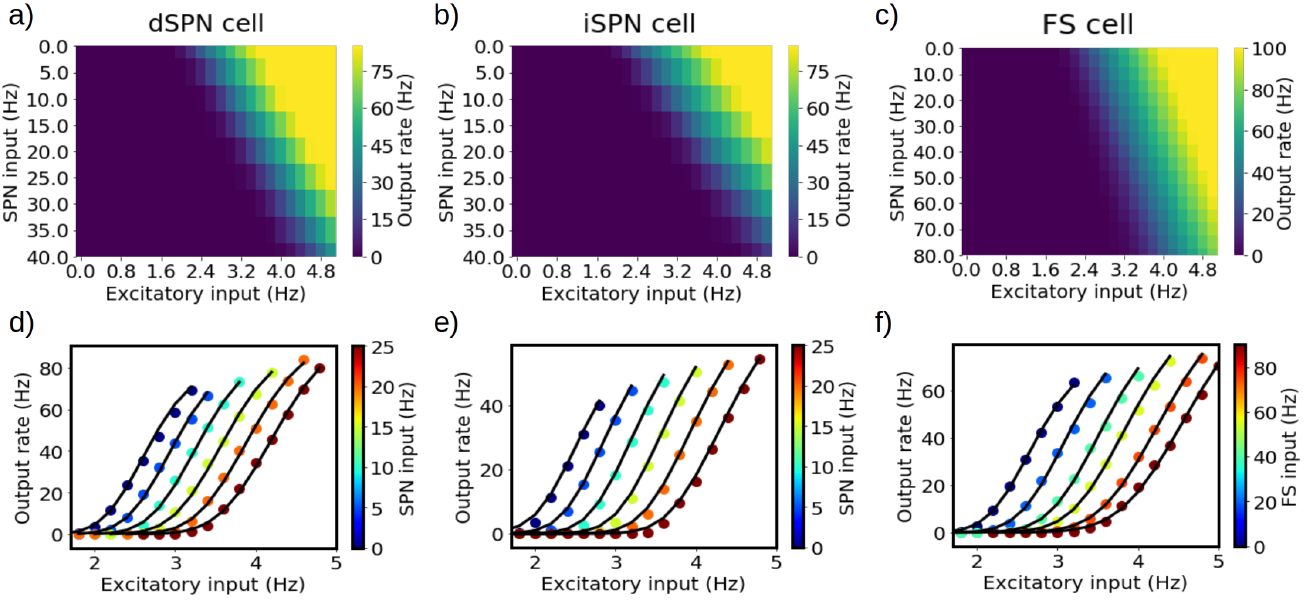
Numerical transfer function (a,b,c) and the corresponding semi-analytical approximation fitted from Eq.5 (d,e,f) for each cell-type. Solid lines in panels d,e,f correspond to the firing rates obtained from Eq.5 while filled-circles correspond to the numerical results. The colormaps in the panels a,b,c indicate the output firing rates obtained from each cell-type for the indicated input. Different circle-colors in panels a,b correspond to different input values of *ν*_*SPN*_, while in panel c correspond to different input values of *ν*_*FS*_. For illustration purposes the panels a,b,d,e correspond to the particular case *ν*_*FS*_ = 0, nevertheless the fit was performed for the full range in the three variables.

### 3.2 *δ, θ, β*, and *γ* rhythms

Oscillations in neuronal activity constitute a key element in brain function, participating in the processing and transmission of information, defining different brain states and being potentially altered in neurodegenerative diseases. As such it’s crucial for the mean-field to properly respond to brain rhythms observed in the corresponding brain area. The striatum exhibits activity in a wide range of oscillatory frequencies, and it is believed that transitions in the spectral power can be of great importance for functions such as motor control and decision making [16]. In Fig.3 we analyze the response of the striatum mean-field to stimulation at the main frequency ranges observed in the striatum. We compare the results of the mean-field with the ones of the spiking-network super-imposed for each case. As we can see the mean-field can correctly reproduce the response of the system for the different frequency bands. For the higher frequencies the instantaneous firing rate of the spiking network becomes more difficult to define, and as the oscillatory frequency becomes closer to characteristic time of the mean-field (see methods), then slight discrepancies may appear (as seen for example in a small phase delay for the mean-field at the high *γ* range).

**Fig. 3.**
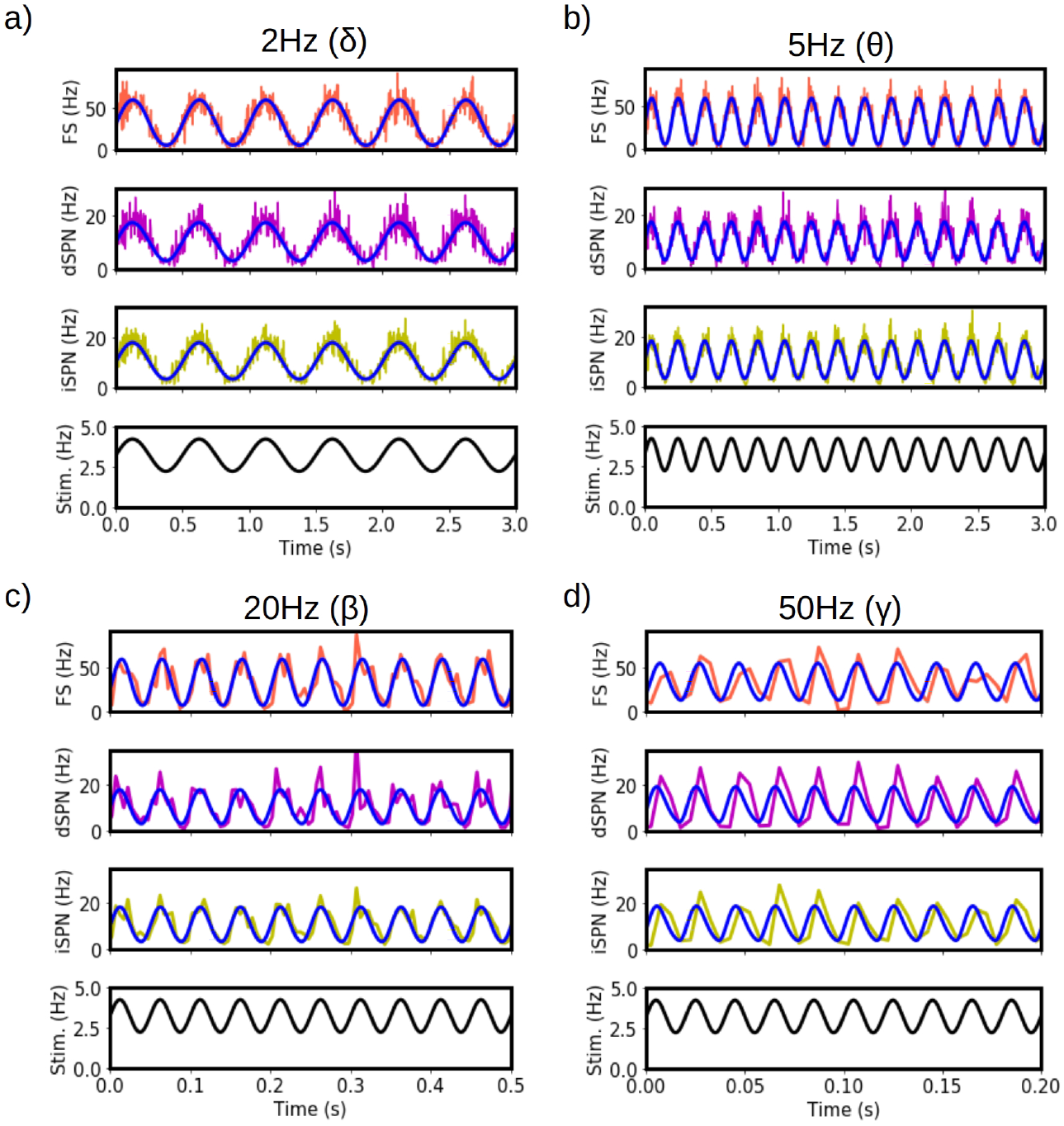
Response of the systems to *δ* (a), *θ* (b), *β* (c), and *γ* (d) rhythms. Results from the meanfield (solid blue lines) are superimposed to the firing rates obtained from the spiking-network model of the striatum.

### 3.3 Dopaminergic effects in striatal neurons

It is well known that dopamine (DA) plays a key role in the regulation of the activity in striatal neurons [4]. It is normally accepted that projections of dopaminergic neurons from the substantia nigra pars compacta (SNc) towards the striatum plays a fundamental role in functions such as motor control and learning, while alterations in the dopaminergic projections are directly related with neurological disorders such as Parkinson’s disease and schizophrenia [17, 18]. Thus, capturing the effect of DA in the striatum with our mean-field model is of fundamental importance.

Striatal neurons exhibit two main types of dopaminergic receptors, D1 and D2. D1 receptors are present in dSPN neurons and generate an increase in dendritic excitability and glutamatergic transmision, while D2 receptors present in iSPN neurons generate the opposite effect with a decrease in the neuronal activity [17, 18]. D1 receptors are known to induce activation of adenylyl cyclase, which increases the level of cytosolic cAMP with the subsequent activation of protein kinase A and which can lead to an increase in the expression of AMPA and NMDA receptors and the enhancement of NMDA mediated currents [19, 20]. On the contrary, D2 receptors are believed to inhibit adenylyl cyclase and it is thought that they act reducing neuronal excitability and neuronal response to glutamatergic inputs [19, 20]. To model these effects of dopamine in dSPN cells we will assume the increase of excitability due to D1 activation in dPSNs can be described as an increase in the glutamatergic conductance (*Q*_*e*_ in our model) together with the action of a depolarizing current, while the activation of D2 receptors in iSPN generates the opposite effect (a decrease in *Q*_*e*_ and a hyperpolarizing current). We show the result in Fig.4 for the implementation of these two effects in our model. In Fig.4.a we show the response of the mean-field for a DA-induced step current with gaussian shape and amplitude +50pA for dSPN’s and -50pA for iSPN’s. In Fig.4.b we show the response of the mean-field for a time-varying quantal conductance of excitatory synapses (*Q*_*e*_) with gaussian shape and maximum amplitude of 10% measured from baseline. Finally in Fig.4.c we show the response for the two effects, both of gaussian shape. In all cases the response of the spiking-neuron is superimposed to the response of the mean-field. We can see that the mean-field can correctly capture the response of the spiking network to the variations on excitabilty.

**Fig. 4.**
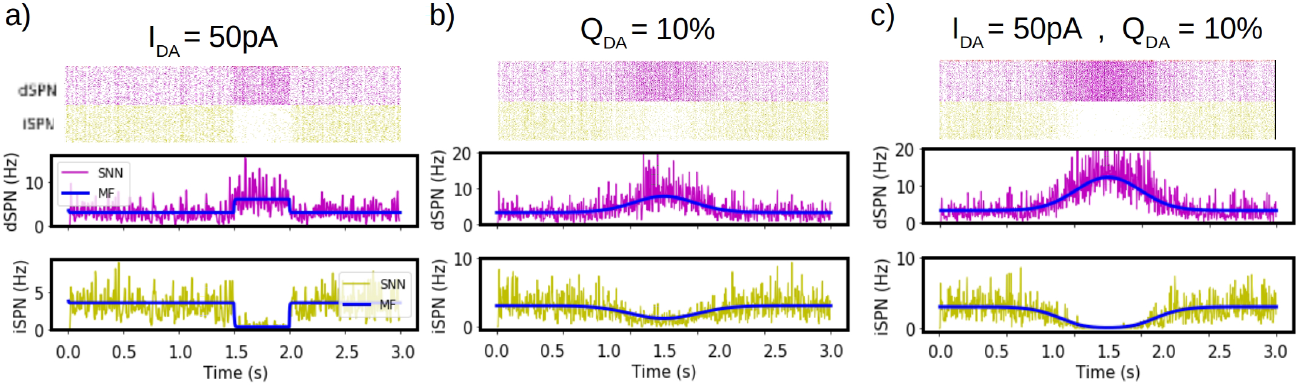
Dopaminergic effects on striatal cells. a) Response of the system to dopaminergic driven current affecting dSPN and iSPN cells. The currents considered excitatory (positive) for dSPN cells exhibiting D1 dopamine (DA) receptors and inhibitory (negative) for iSPN cells exhibiting D2 receptors. We show the raster plot obtained from network simulations and the corresponding mean firing rates obtained from the network and from the mean-field for dSPN and iSPN cells. The dopaminergic current is simulated as a square function. b) Response of the system to a transient change of synaptic conductance. As in panel (a), the change is considered positive for dSPN cells and negative for iSPN cells. In this case the variation in conductance is modeled as a gaussian function on time. c) Response of the system to a combined DA current and conductance variation accounting for the multiple effects of dopamine in SPN cells.

### 3.4 Reinforcement learning

It is known that basal ganglia play a key role in the processes of learning and decision making [1, 2, 21]. The standard vision is that the dopaminergic pathways, involving the projection of DA neurons from Substantia Nigra pars compacta (SNc) to the striatum, work as reward signal within a reward-learning mechanism in the brain. The release of dopamine by SNc projections in response to an action regulates the strength of the local synaptic coupling on the corticostriatal circuit, modulating the output signal of the striatum (via the ‘direct’ and ‘indirect’ pathways) and thus modifying the response/action to a certain stimulus [17, 18, 21]. In the direct pathway, activation of dSPN cells due to cortical input projects inhibitory signals to the SNr and to the internal segment of globus pallidus (GPi). The inhibition of GPi/SNr generates a disinhibition of thalamic glutamatergic neurons, which project to the cortex. This promotes the initiation of locomotor activity. On the contrary, in the indirect pathway, activation of iSPN cells due to cortical input generates a disinhibition of the SNr and GPi (via the external segment of globus pallidus and the subthalamic nucleus), which in turn inhibits the thalamic glutamatergic neurons, diminishing locomotion activity [17, 18, 21]. Thus, in a simplified manner, higher activity in dSPN cells induces the realization of an action, while higher iSPN activity inhibits the action in a go/no-go scenario [18, 22]. It’s known that DA activation of D1 receptors present in dSPN cells tends to induce Long Term Potentiation, while activation of D2 receptors tend to induce Long Term synaptic Depression in iSPN [18, 19], modifying the activation of the direct/indirect pathways. This DA modulation of the synaptic strength between cortical input and dSPN/iSPN cells (controlling the direct/indirect pathways) is a well accepted vision on the mechanism of Reinforcement Learning (RL) in the brain. In this section we show how a basic model of RL can be implemented within our mean-field model of the striatum. To this end we will consider a simple task where two stimuli (Stim. A and Stim.B) are presented and a binary (go/no-go) decision has to be made. A diagram of the model is shown in Fig.5.a. Each stimulus is represented by a different cortical input (*ν*_*e*−*A*_, *ν*_*e*−*B*_) to a single functional domain within the striatum (represented by a single mean-field model). A decision towards Action-1 (go) is made with probability given by:

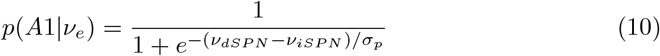

where *σ*_*p*_ is a constant. The probability of taking Action-2 is given by *p*(*A*2|*ν*_*e*_) = 1 − *p*(*A*1 |*ν*_*e*_). Given the decision taken by the subject a reward/punishment signal is applied represented by a transient increase or decrease of the dopaminergic level at the striatum. We will assume that Stimulus A is associated with a reward signal for the choice of Action-1 (and a punishment for Action-2), and the opposite for Stimulus B. The dopaminergic signal can modulate the strength of the corticostriatal synaptic coupling for each cortical projection. For simplicity we will assume that the dopaminergic signal is instantaneously applied during the stimulation period and that the modification of synaptic coupling by DA occurs in a heterosynaptic way, where the synaptic changes are independent of the post-synaptic activity. The synaptic rule used in our model is given by:

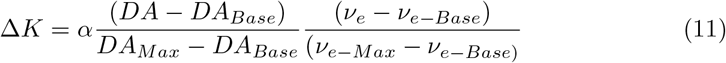

where Δ*K* is the change in synaptic strength (i.e. change in synaptic convergence, see Eqs. 7, 8 and 9), *DA*_*Base*_ (*ν*_*e*−*Base*_) is the baseline level of dopamine (cortical activity), *DA*_*Max*_ (*ν*_*e*−*Max*_) is the maximum level of dopamine (cortical activity), and where *α* is constant. Following the fact that activation of D1 receptors present in dSPN cells tend to induce Long Term Potentiation, while activation of D2 receptors tend to induce Long Term synaptic Depression in iSPN, we will take *α* = +*/* − 0.2 with the positive sign for cortico-dPSN synapses and the negative sign for cortico-iSPN synapses. Thus, according to this rule, the increase (decrease) of the dopaminergic level correlated with the activation of a specific cortical input generates a potentiation (depression) of the cortico-dSPN synaptic coupling, while the opposite effect is caused in the cortico-iSPN coupling.

We show the results of the simulations in Fig. 5. In Fig.5.b we show the time-series of the firing rates and stimulus application during the simulation. In the stimulation protocol the stimulus A and B are presented alternately (indicated with the cyan and magenta shaded bars in the right column). In Fig.5.c we show the decision taken after each stimulus presentation. We see that at the beginning of the simulation the subject picks randomly between Action 1 and 2. As the simulation progresses the subject modifies its choice according to the training and as a result of the preferential activation of dSPN and iSPN cells to each stimulus.

**Fig. 5.**
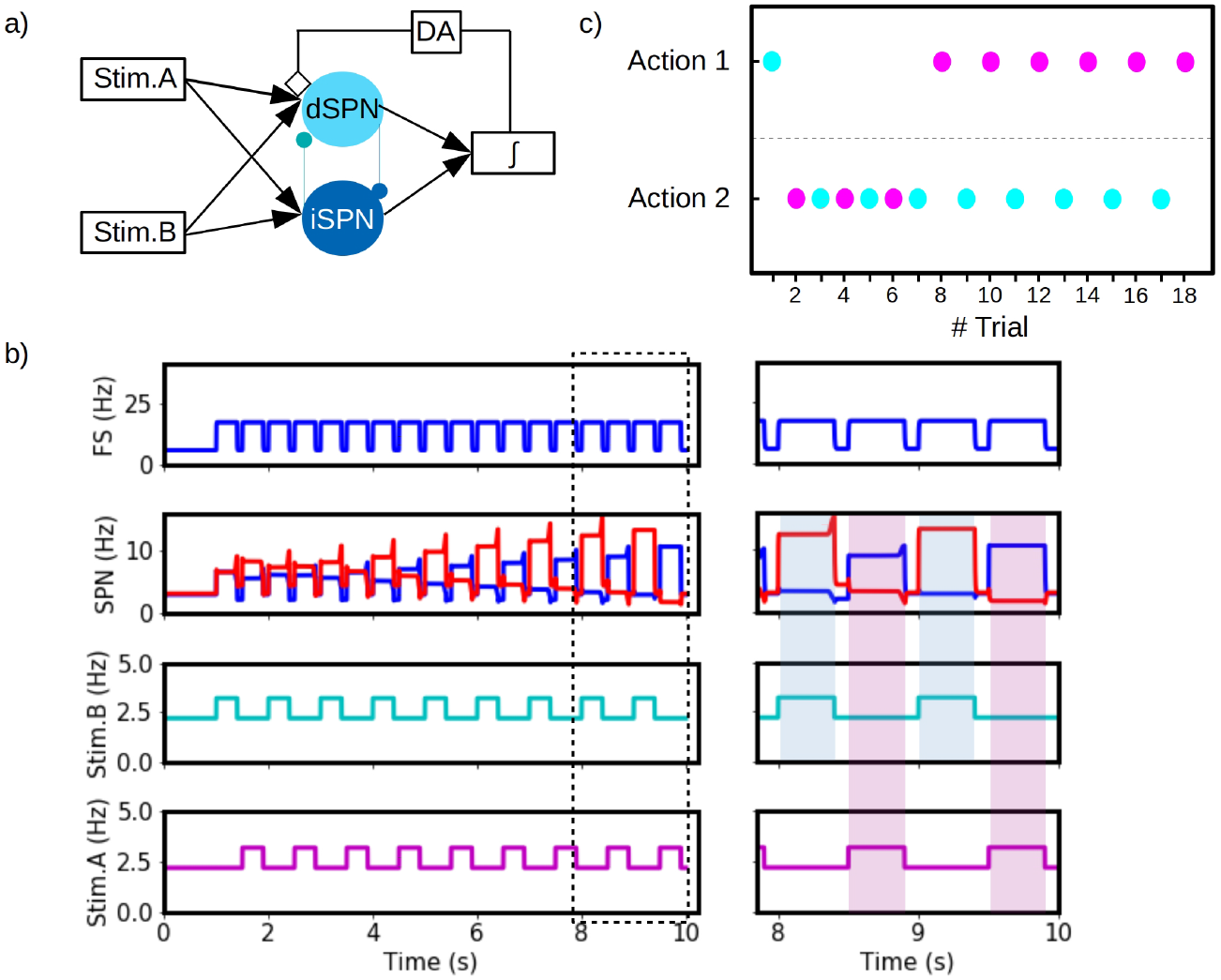
Basic implementation of reinforcement learning (RL) with the mean-field model of the striatal microcircuit. a) Diagram of the RL model. Two different sensory stimulus (A and B) are represented by two independent cortical projections towards the striatum. The output of dSPN and iSPN cells are used to decide between two possible actions (Action 1 and Action 2) according to Eq.10. The resulting action will generate a reward or punishment dopaminergic signal (DA) which can modify corticostriatal synaptic weights according to Eq.11. b) Simulation of the RL process. We show the cortical activity (Stim. A and Stim.B, bottom panel), the dSPN, iSPN (red and blue lines respectively, second plot from the top) and the FS response. On the right column we show the detailed for timeperiod indicated with the squared area (between 8-to-10 seconds approximately). We can see that by the end the simulations the dSPN and iSPN cells respond selectively to each stimulus type. c) Action selection for each stimulus presentation. During the first trials the system is incapable of distinguishing the correct action to take under each stimulus type. By the end of the simulation, the system can correctly select the Action 1 for Stim.A (magenta circles) and Action 2 for Stim.B (cyan circles).

## 4 Discussion

In this paper we have presented a multiscale investigation of the striatum microcircuit. We have built the model based on a recently introduced formalism that follows a bottom-up approach, starting at the single-cell and (spiking) network levels, which allowed us to incorporate cellular and synaptic specifities of striatum within the mean-field formulation. The single-cell parameters were obtained from fitting of AdEx neuronal model to electrophysiological data of striatal projection neurons (dSPN, iSPN cells) and fast-spiking interneurons (FS cells), and synaptic connectivity information was extracted from experimental and simulated data. We have tested the models by analyzing their response to different oscillatory rhythms found in the striatum and we have validated the results by systematic comparison of the mean-field model with the corresponding spiking network model. We have shown that the mean-field can correctly reproduce the results of the spiking-network for the main oscillatory patterns observed in the striatum (*δ, θ, β*, and *γ*).

We have also studied the effect of dopamine on striatal neurons. We have shown how the action of dopamine can be included within the mean-field model and we have validated our results by comparison with the results of spiking networks. This analysis allowed us to show how modifications at the cellular level can be incorporated within the mean-field model which can in turn predict and capture the emergent changes at the network level generated by them, and in addition has provided further validation to our model. Alterations in dopaminergic levels are in the center of several neurological disorders such as Parkinson’s disease and schizophrenia [17, 18], for which the capacity of the model to capture its effects is of great importance. This analysis can be further extended by incorporating other BG nuclei following a similar approach, which should be done in future studies.

Furthermore, we have presented a basic implementation of reinforcement learning (RL) with the mean-field model, which constitute one of the best known functions of the striatum. Models of reinforcement learning considering the striatum have been studied for decades, and the role of dopaminergic projection onto striatal cells is nowadays accepted as one of the main mechanisms of RL in the brain [17, 22]. We have proposed a model with a simple task with two stimuli and two possible actions, within a basic circuit for reward/punishment based on dopaminergic action on striatal cells. Our model can correctly capture the learning process which shows the capabilities of the model to reproduce specific brain functions and in particular its potential implementation for modelling behavioural experiments. The RL model proposed can very easily be improved to capture more biologically complex scenarios by introducing, for example, the dopaminergic effect on SPN firing rates, adding other brain circuits participating in the RL process and proposing more complex tasks.

It is important to note that methods to estimate brain signals (LFP, EEG, MEG, fMRI) from this type of mean-field models have been recently developed [23, 24]. While such methods were not used here, they open the possibility to directly compare the striatal model to experimental recordings.

Finally, our model is suitable to be incorporated in large-scale simulators such as the The Virtual Brain (TVB) platform [25], as shown previously for cerebral cortex, where different brain states and their responsiveness could be simulated at the wholebrain level [7, 26]. Because the implementations follow the same formalism, the present model can be easily incorporated in TVB simulations, leading to an integrated cortex - basal ganglia model at the whole-brain level. This will also constitute a next step for future investigation.

